# DALE-Eval: A comprehensive cell type-specific expression deconvolution benchmark for transcriptomics data

**DOI:** 10.1101/2025.07.31.667984

**Authors:** Mengying Hu, Martin Jinye Zhang, Maria Chikina

## Abstract

Deconvolution of bulk transcriptomic data unlocks rich, cell type–specific insights from complex tissue samples. While cell type fraction deconvolution has been extensively developed and bench-marked, the next frontier—cell type-specific expression (CTSE) deconvolution—remains largely underexplored. Here, we introduce DALE-Eval, a comprehensive benchmark of CTSE methods, featuring novel evaluation metrics, identification of key drivers of performance, and practical recommendations for maximizing the value of CTSE in downstream applications.

## Main

Computational deconvolution of bulk transcriptomic data provides cell type-resolved insights into heterogeneous mixtures by disentangling signals arising from overlapping gene expression across different cell types. Significant methodological developments and benchmarking efforts in this area[1] have predominantly focused on estimating cell-type proportions (fraction estimation), with pioneering methods demonstrating substantial promise, such as BayesPrism[2], MuSiC[3] and CIBERSORTx[4]. However, beyond fraction estimation, recent advances now allow for more refined deconvolution through cell type-specific expression (CTSE), which estimates expression for each cell type present in bulk data without requiring single-cell profiling[2, 4–9]. This approach holds substantial potential for translational studies when single-cell data are unavailable or impractical, facilitating downstream analyses such as cell type-specific eQTL mapping, differential expression analysis, and unsupervised clustering[10–12]

However, the extent to which CTSE deconvolution is truly reliable remains unclear. Previous evaluations[5–8, 13] have been limited in scope—focusing on a small number of methods, restricting to specific tissues, and omitting comparisons to simple but informative baselines, such as bulk or fraction-adjusted bulk expression. Key questions remain unresolved: Do current methods offer real gains over naive baselines? What determines deconvolution accuracy? And are the estimated profiles truly capturing cell type–specific expression variation, or do they reflect cross-cell type contamination? Clarifying these issues is essential for robust downstream applications.

To address these gaps, we introduce DALE-Eval—the first comprehensive framework for benchmarking CTSE deconvolution (Figure 1). Using multi-individual single-cell RNA-seq data from PBMC, brain, and tumor, we generate realistic test sets with known ground truth (Extended Data Table 1). We evaluate leading methods including CIBERSROTx[4], InstaPrism/BayesPrism[2, 14], bMIND[5], EPICunmix[6], Unico[7], ENIGMA[8], and TCA[9], alongside baseline approaches (Methods; Extended Data Figure 1), using independent data for reference construction mirroring real world applications (Supplementary Note 1). Our evaluation emphasizes inter-sample gene-level correlations (Supplementary Note 2) and introduces novel assessments of cell type specificity (Supplementary Note 3), stratifying performance across a broader range of evaluation scenarios.

**Figure 1:**
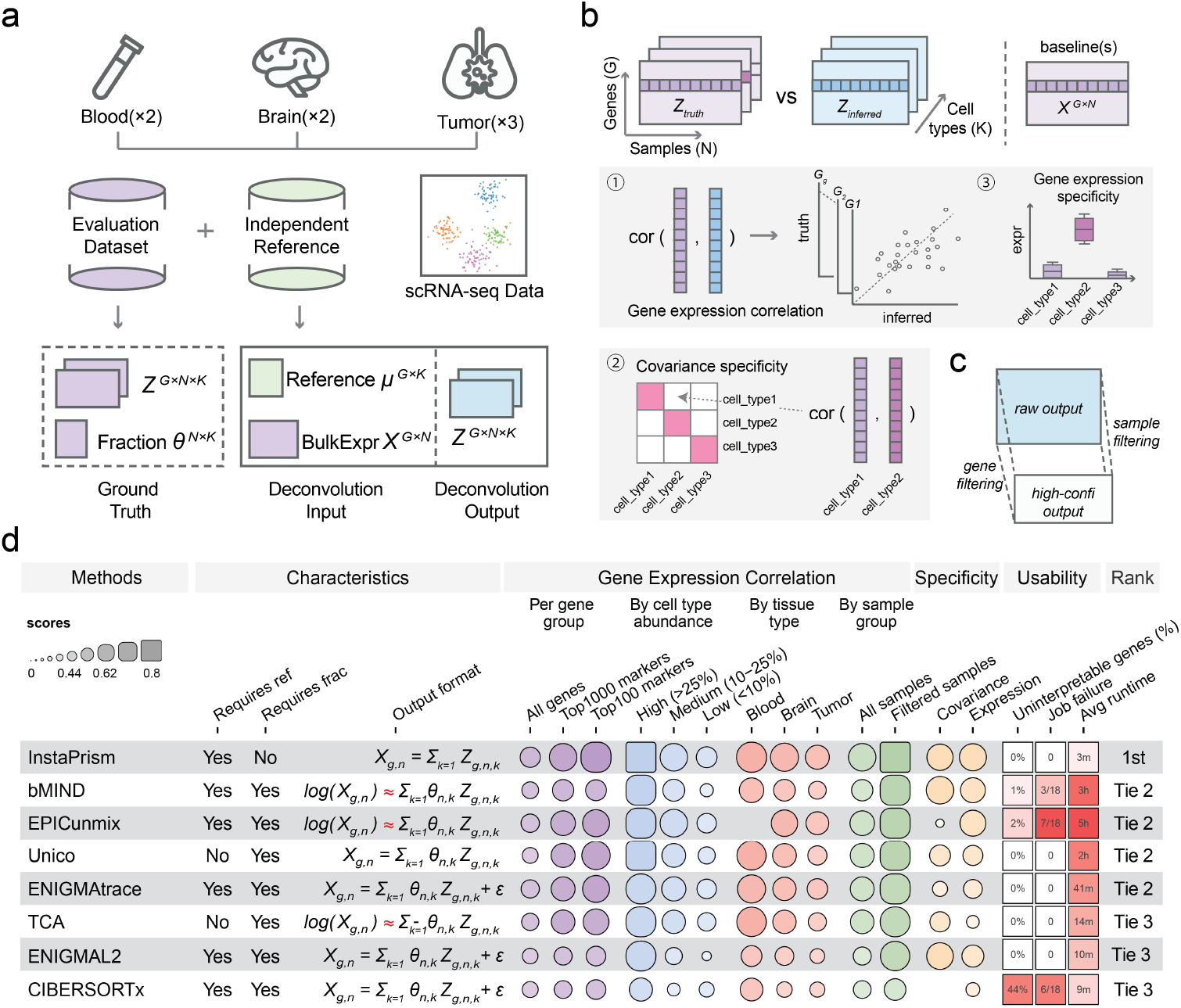
Overview of DALE-Eval workflow and summary of the benchmarked methods. **a)** Benchmarking data construction using scRNA-seq data. **b)** Evaluation metrics and baselines. Deconvolution performance was assessed by (1) inter-sample gene correlations (computed for each gene-cell type pair across samples), (2) covariance and (3) expression specificity. Bulk and fraction-regressed bulk data are used as baselines. **c)** Proposed reliable CTSE subset: Raw CTSE profiles should be filtered by gene specificity and by samples with sufficient cell type proportion to ensure reliability. **d)** Performance summary. Gene correlation and specificity scores (0–1; higher is better) are summarized across all datasets tested.

Our benchmarking results revealed four critical insights. First, when averaging performance across all genes, all methods performed poorly in recovering true inter-sample variations in most cell types (Figure 2a; Extended Data Figure 2), with the best-performing method reaching a mean correlation of only 0.4. In fact, under this context, most methods did not even outperform bulk baseline, or provided only minimal improvement (correlation difference < 0.05).

**Figure 2:**
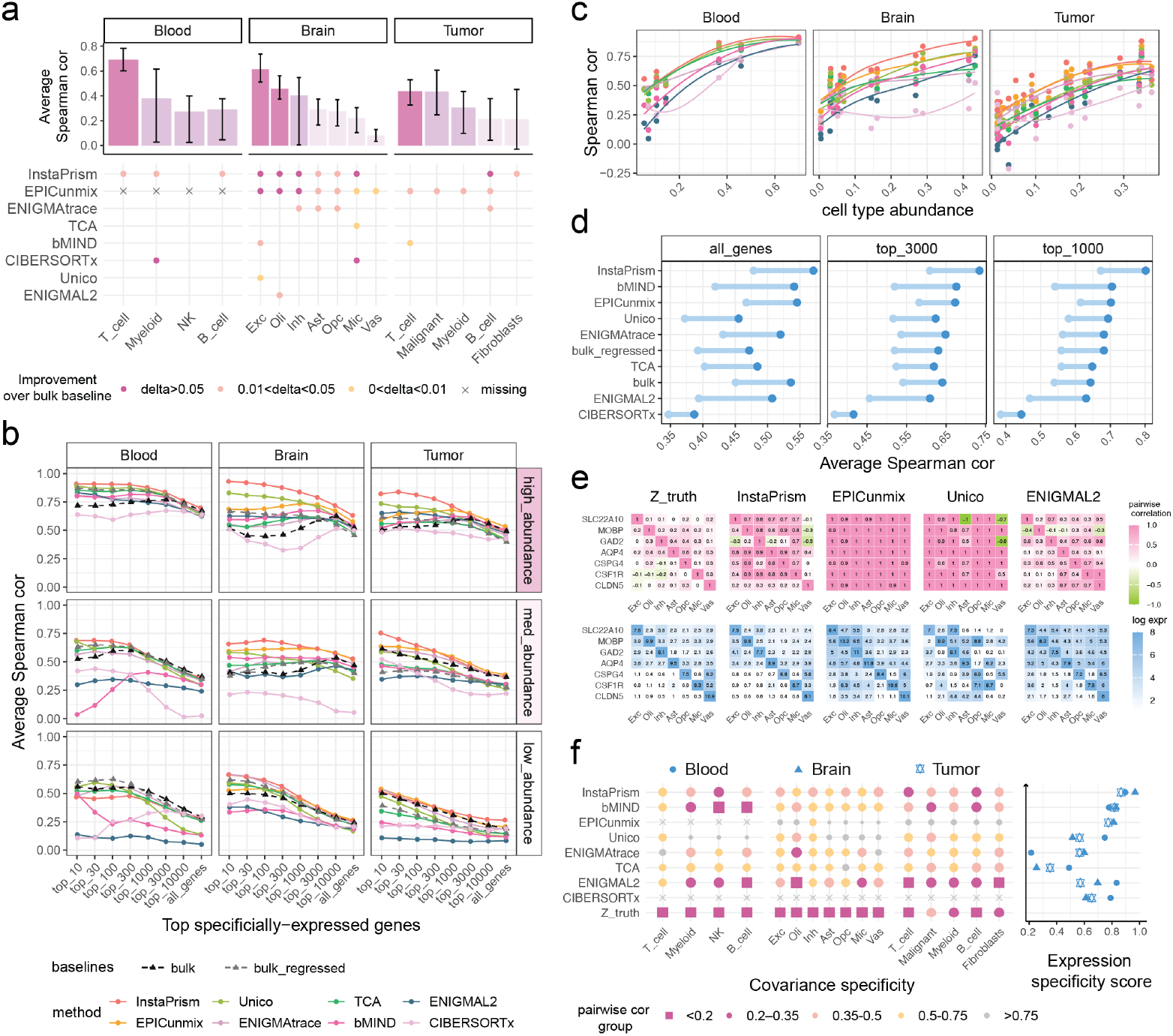
Evaluation of CTSE deconvolution methods. **a)** Deconvolution accuracy averaged across all genes. Top: mean performance per cell type, with error bars denoting the min and max across methods. Bottom: gene correlation improvement over bulk (non-improving omitted); cell types ordered and color-coded by abundance: high (≥ 0.25), medium (0.1–0.25), and low (< 0.1). **b)** Average inter-sample correlation of cell type-specific genes. Line plot of mean prediction accuracy across the top *n* specifically expressed genes, stratified by cell type abundance groups. **c)** Prediction accuracy vs. cell-type abundance: each point shows the mean correlation across the top 1,000 marker genes. **d)** Dumbbell plot of mean gene correlation before (light blue) and after (dark blue) filtering out samples with cell-type fraction < 0.1. **e)** Illustration of marker gene specificity in brain cell types. Top (covariance specificity, pink): Pairwise correlation of each marker gene’s expression in its target cell type versus every other cell type. (diagonal element represent target gene-cell type pairs.) Bottom (expression specificity, blue): Mean expression of marker genes across all cell types. Example shown for ROSMAP_AD92_Xiong2023 dataset. **f)** Summary of covariance and expression specificity. Lower pairwise correlation and higher expression specificity scores indicate better cell type specificity. Analyses are based on the top 30 marker genes per cell type.

Second, we investigated the performance across gene groups with various levels of specificity. Specifically expressed genes, or marker genes, were defined as those with elevated expression in one cell type relative to others, derived and ranked using reference data (Methods). Using this stratification, we observed that prediction accuracy was generally highest for cell type-specific genes and declined progressively for less specific ones (Figure 2b; Extended Data Figure 3). This suggests that gene specificity can serve as valuable prior knowledge to prioritize genes that are more likely to be accurately deconvolved. However, we caution that some methods (e.g., bMIND and EPICunmix) deviated from this pattern and performed poorly on the strongest marker genes in some cell types, underscoring the need for extra configuration for gene-prioritization. Additionally, we found that the benefit of gene stratification varied across cell type abundance groups and tissue types. For blood, the simple fraction-regressed bulk baseline already effectively captured variation among top-specific genes, and few methods outperformed it. However, for brain and tumor, deconvolution indeed provided additional gains in expression estimation, particularly for abundant cell types.

Third, we examined the impact of cell type abundance. As expected, abundance differences explained much of the variation in prediction performance, with more abundant cell types yielding more reliable estimates across all methods (Figure 2c). This also uncovers a hidden insight: inter-sample variation may be better preserved within samples with higher cell-type abundance. To test this, we recalculated gene correlations using only samples with cell type fractions above 0.1. Our results suggested that this filtering substantially improved deconvolution accuracy (Figure 2d; Extended Data Figure 4), likely due to removal of poorly represented expression estimates from low-abundance samples. However, we note that the 0.1 threshold is only a rough guideline and there is a trade-off between number of samples retained and the choice of this threshold. In practice, since cell-type abundance still varies after filtering, the number of genes considered reliable for prioritization should be scaled accordingly, for example, selecting more genes for abundant cell types and applying more conservative criteria for rare ones.

Fourth, we assessed the cell type specificity of the estimates. In principle, each cell type represents an independent transcriptional unit; gene expression variation in one cell type (e.g., *BCL6* expression in B cells) does not imply similar variation in another (e.g., T cells). This biological independence is supported by the minimal cell type co-variation observed in ground truth profiles (Figure 2e). However, when summarizing covariance structures in deconvolved profiles (Methods), we identified a concerning leakage of expression signals across all tested methods, reflected by elevated pairwise correlations between cell types (Figure 2e,f). This lack of covariance specificity was observed not only for cell type marker genes, but for all genes in general (Supplementary Note 3).

We reasoned that the suspiciously high co-variation structure points to two possibilities: **(1)** the observed variation originates from true biological differences in just one dominant cell type, but this signal is mistakenly propagated to others due to the decomposition process. This scenario is common when the bulk expression is heavily dominated by a major cell type or when analyzing strong marker genes—both cases in which deconvolution performance tends to be highest; and **(2)** the inferred expression represents a proportion adjusted cell type average and thus does not yield meaningful cell type–specific signals. The second scenario likely applies to the majority of estimated values; it also explains why when considering the overall gene correlation, performance is only moderately improved or no better than the bulk baseline.

As such, we caution that while most methods output a complete gene-by-sample-by-cell type CTSE tensor, many entries in this tensor are unreliable. Prioritizing top cell type–specific genes using external reference data offers a partial solution (Figure 2b), however, a rigorous framework is still needed to assess confidence in each gene’s estimates. One potentially valuable resource is the CTSE profiles themselves. By examining expression specificity (Methods), we observed that some methods accurately captured the relative expression patterns—for example, correctly identifying *AQP4* as predominantly expressed in astrocytes but not in others (Figure 2e). *Post hoc* analysis of these profiles may therefore guide the selection of genes that are truly cell type-specific and whose variation likely reflect true cross-sample differences (Supplementary Note 4).

CTSE deconvolution remains challenging: most methods offer limited gains over naive baselines and often fail to produce truly cell type–specific profiles. Nonetheless, our results show that accurate deconvolution is feasible for a subset of genes—particularly those with high cell type specificity and in abundant cell types. Substantial improvements will depend on fundamentally rethinking current approaches, including better use of covariance structure in single-cell references and rigorous uncertainty quantification to flag unreliable estimates. Our framework lays the foundation for these advances and provides practical guidance for real-world applications.

## Methods

### Single-cell RNA seq datasets

We collected single-cell RNA-seq data from human samples for this benchmark using the following selection criteria: (1) for benchmarking datasets, we selected large-scale single cell studies containing more than 30 individual samples; (2) for reference datasets, we chose scRNA-seq dataset from a matched tissue with detailed cell type annotations at both cell-type and sub–cell-type (cell-state) resolution. A detailed description of the datasets used in this benchmark is provided in Extended Data Table 1

For each dataset, raw gene expression count matrices were downloaded and processed for quality control. Cells with fewer than 500 detected genes, mitochondrial transcript proportions exceeding 20%, or total UMI counts below 1,000 were excluded. Genes detected in fewer than 100 cells were also removed. The original cell type annotations were used as cell type identifiers.

### Benchmarking data

Using the scRNA-seq datasets collected, we established benchmarking data with the following components: (1) ground truth cell type–specific expression (CTSE) profiles *Z ∈* ℝ^*G×N ×K*^, (2) ground truth cell type fractions *θ* ∈ ℝ^*N ×N*^, and (3) bulk gene expression profiles *X* ∈ ℝ^*G×N*^ used as input for deconvolution.

Specifically, the count data were first normalized to counts per million (CPM), then averaged by their source sampleID and cell type labels. For each cell type *k*, its specific expression *Z*_,,*k*_ ∈ ℝ^*g×n*^ was computed as the averaged gene expression of cells from the same sampleID for this cell type. For each sample *j*, the cell type fraction *θ*_*j*_ *∈* ℝ^1*×K*^ was computed as the proportion of cells from each cell type in the single cell data, and the bulk expression *X*_*j*_ ∈ ℝ^*G×*1^ was calculated as the average gene expression of all cells within that sample. For cell types with zero fraction, the corresponding CTSE values were set to zero. A minimum of 10 cells per cell type per sample was required to construct a CTSE profile; otherwise, that cell type will be discarded for that sample.

The final components in the benchmarking data satisfy the following relationship, which is also consistent with the assumption of the deconvolution model:

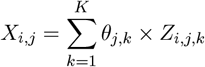

We note that the choice of generating pseudobulk samples by aggregating cells from their true biological samples helps preserving the natural heterogeneity of CTSE profiles. In contrast, simulating additional “bulk” samples—for example, generating 100 samples from scRNA-seq data with only 10 true samples—may result in less biological variability[1, 15]. and inevitably involves additional technical choices, such as sampling methods, distributional assumptions, and parameter settings, which may confound benchmarking results.

### Benchmarking framework

For each benchmarking scRNA-seq dataset, 80% of samples (by sample ID) were randomly assigned to generate the benchmarking data, while the remaining 20% were used to construct an internal reference for complementary analyses assessing the impact of reference selection (Supplementary Note 1). Note that sample split was performed at the sample level rather than by cell type to prevent information leakage that would occur if cells from the same individual appeared in both the test and reference sets.

Unless otherwise specified, all reference data used for deconvolution in this study were derived from independent, tissue-matched scRNA-seq datasets. This emulates realistic scenarios in which the cell type composition of the bulk data is unknown and researchers must rely on external, tissue-matched single-cell references to guide deconvolution analysis[1].

For reference-based methods, each algorithm received its required reference input constructed from the reference scRNA-seq datasets (see “Reference construction” for details). For methods requiring cell type fraction as input, we supplied fractions estimates inferred by InstaPrism/BayePrism, which used the same scRNA-seq dataset as the reference-based methods. This approach mimics real-world scenarios where ground-truth fractions are unavailable; InstaPrism/BayesPrism was chosen due to its superior fraction estimation accuracy demonstrated in multiple independent benchmarking studies[15, 16].

### Reference construction

Each reference scRNA-seq dataset was processed as follows to generate the necessary reference input for each deconvolution method, in line with the guidelines for each method.:

#### bMIND/EPICunmix reference

While bMIND accepts both a mean prior and a covariance prior, CTSE estimates are only available for genes with positive definite covariance matrices when the covariance prior is provided, leading to substantial missing values. To ensure completeness and comparability with other methods, we supplied only the mean prior, calculated using the `get_prior()` function from `EPICunmix` R package. EPICunmix builds directly on bMIND results, so its reference construction procedure is considered the same as described above.

#### CIBERSORTx reference

CIBERSORTx reference matrices were constructed using the `build_model_cibersortx()` function from the `omnideconv` R package[17]. To manage computational load, datasets exceeding 400,000 cells were downsampled to around 50,000 cells and only the 10,000 most variable genes were supplied. Note that while CIBERSORTx reference typically contained around 3,000 genes, the final CTSE output always returns the full bulk gene set (the same number of genes as in the input bulk data).

#### ENIGMA reference

ENIGMA requires a mean prior matrix with dimensions *G × K*. We constructed this matrix by averaging CPM-normalized single-cell expression values within each cell type, then supplied it as the reference input to both optimization modes, ENIGMAL2 and ENIGMAtrace.

#### InstaPrism reference

We built InstaPrism reference using the `refPrepare()` function from `InstaPrism` R package, which takes single-cell expression data along with cell type and cell state labels as input. For cell state labels, we used the refined annotations provided in the reference datasets (e.g., subtypes or sub-cluster labels that offer greater granularity than broad cell types). For malignant populations, we followed the developers’ recommendation to use sample identifiers as cell state labels[2].

### Implementation of deconvolution methods

#### bMIND

bMIND[5] was implemented using the `MIND` R package (version 0.3.3), with three required inputs: log-transformed bulk expression, the mean prior (as described above), and InstaPrism-inferred cell type fractions.

#### CIBERSORTx

CIBERSORTx[4] was run within a local singularity environment using the “High Resolution” mode, following the guidelines provided at https://open.bioqueue.org/home/knowledge/showKnowledge/sig/cibersortx-hires. Three required inputs were supplied: bulk expression, the CIBERSORTx signature matrix (as described above), and InstaPrism-inferred cell type fractions.

#### ENIGMA

ENIGMA[8] was implemented using the `ENIMGA` R package (version 0.1.6) with default settings; the trace norm and maximum L2 norm models are referred to as ENIGMAtrace and ENIGMAL2 respectively. Three required inputs were supplied for both models: bulk expression, the ENIGMA reference matrix (as described above), and InstaPrism-inferred cell type fractions. By default, ENIGMA applies further normalization to the raw deconvolution results. In the Extended Data, we also evaluated the unnormalized outputs, referred to as ENIGMAtrace(raw) and ENIGMAL2(raw).

#### EPICunmix

EPICunmix[6] was implemented using the `EPICunmix` R package (version 0.0.1). It directly takes the posterior estimates from bMIND, along with log-transformed bulk expression and cell type fractions inferred by InstaPrism, as input. Notably, although the `EPICunmix` package provides its own implementation of bMIND via `EPICunmix::bMIND()`, we did not implement this function because its output for marker genes in their target cell types is largely constant. (data not shown).

#### InstaPrism/BayesPrism

InstaPrism[14] was implemented using the `InstaPrism` R package (version 0.1.6). This method provides a faster implementation of BayesPrism[2] while maintaining identical outputs. Two inputs are provided as required: the bulk expression data and reference profiles (as described above).

Notably, InstaPrism produces a unique CTSE profile *Z* where the bulk expression is represented as a direct linear combination of 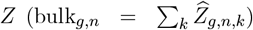. For each sample, the sum of gene expression values for each cell type corresponds to its relative fraction. As such, gene expression values in *Z*_*g,·,k*_ are not directly comparable across samples, since each sample has a different library size in *Z* and must therefore be normalized. For example, high gene expression values do not necessarily indicate higher cell type-specific expression, but may simply reflect higher cell type fractions in that sample.

To enable inter-sample comparisons, gene expression values in *Z* were normalized by dividing each cell type’s values by a scaling factor based on its total expression in each sample. To avoid inflation when a cell type’s fraction was very small, a small constant was added to both the numerator and denominator:

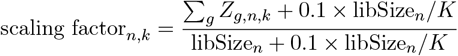

where libSize_*n*_ is the total library size for sample *n* and *K* is the number of cell types.

#### TCA

TCA[9] was implemented using the `TCA` R package (version 1.2.1). Although TCA was originally developed for cell type-specific methylation data, it is also applicable to gene expression data and has been included for comparison in several CTSE deconvolution studies[7, 8, 13]. Two inputs were provided: log-transformed bulk expression and cell type fractions inferred by InstaPrism. Genes exhibiting inter-sample variance under 10^−8^ in the log-transformed bulk expression data was discarded, in accordance with the method’s specifications.

#### Unico

Unico[7] was implemented using the `Unico` R package (version 0.1.0), with two required inputs: bulk expression and cell type fractions inferred by InstaPrism. Genes with inter-sample variance less than 10^−4^ in the bulk expression data was discarded, in accordance with the method’s specifications.

### Evaluation metrics

CTSE deconvolution performance was assessed by comparing the inferred cell type-specific expression (CTSE) profiles, *Z*_inferred_, to the ground truth, *Z*_truth_, where *Z* ∈ ℝ^*G×N ×K*^. Note that since the benchmarking and reference datasets come from different scRNA-seq sources and may have nonidentical gene sets and cell-type definitions, we limited all comparisons to overlapping genes and matched cell types only. Performance was quantified using the following metrics:

#### Gene expression correlation

For each matched cell type *k* ∈ {1, …, *K*} and each gene *g* ∈ {1, …, *G*}, we computed Spearman correlation coefficients between the inferred and ground-truth expression vectors across samples, which yields a *G ×K* matrix of correlation values for each method. Samples with a ground-truth fraction of zero (all-zero profiles) were excluded during gene correlation calculation.

We chose Spearman over Pearson correlation as our primary metric because several methods produce log-scaled CTSE profiles 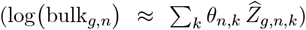 that do not align linearly with either the raw or log-transformed ground-truth expression. However, we also note that evaluation results using Pearson correlation closely match those obtained with Spearman correlation (Extended Data Figure 2).

#### Covariance specificity

We define *covariance specificity* as a measure of how much a gene’s variation is specific to one cell type but not others. To quantify this, for each gene *i*, we computed the pairwise Pearson correlation between its cell type-specific expression profiles across samples. This yields a *G × K × K* array of covariance array *C*, where each entry 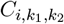 measures how the variation of gene *i* in cell type *k*_1_ is associated with that in cell type *k*_2_:

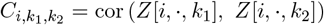

However, not all pairwise correlations in *C* are biologically meaningful. For example, for a B cell marker gene such as *CD79A*, the relevant information is how its expression variation in other cell types covaries with that in B cells, rather than its covariation between unrelated cell types (e.g., T cells and macrophages). Therefore, from the full covariance array *C*, we further summarized the pairwise correlation patterns as follows: for each cell type *k* with marker gene set ℳ_*k*_, we extract

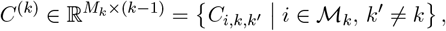

which yields a set of correlation values reflecting the similarity of marker gene expression between its target cell type *k* and all other cell types, with lower pairwise correlation indicating higher covariance specificity.

We note that CIBERSORTx is a special case here, as its CTSE output is largely composed of constant values of 1 and co-variation between different cell types is mostly undefined; therefore we do not consider covariance specificity for this method.

#### Expression specificity

We define *expression specificity* as the degree to which a gene maintains its characteristic expression pattern in *Z*_inferred_ relative to *Z*_truth_—for example, a gene being highly expressed in its target cell type and lower expressed in other cell types.

To quantify this, we first computed cell type-specific expression pattern from *Z*. Specifically, for a given gene *g* we calculated the its average expression in each cell type *k* as:

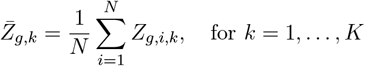

where 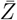 matrix with dimension *G × K*. We then assessed the agreement between 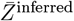 with 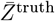 using Lin’s Concordance Correlation Coefficient (CCC)[18], which captures both correlation and their deviation from the line of perfect concordance (*y* = *x*). To stabilize variance, 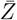 was transformed to log scale prior to CCC calculation.

Specifically, given 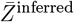 and 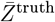 (both already in log scale), for each gene *g*, we computed:

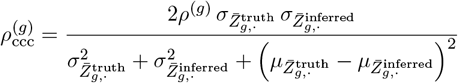

where *ρ*^(*g*)^ is the Pearson correlation coefficient between the vectors 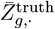 and 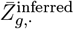, and *µ* and *σ*^2^ denote the mean and variance for each 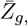_,*·*_ vector respectively. Higher CCC values indicate better expression specificity.

### Evaluation baselines

For this benchmark, we used two simple baselines to assess the information gain from deconvolution: plain bulk and fraction-regressed bulk, both represented as *G× N* matrices without cell type-specific information. Plain bulk baseline corresponds to the original bulk input for deconvolution process, and the fraction-regressed bulk baseline is generated by removing the component of gene expression explained by cell-type fractions. Specifically, for each gene, bulk expression across samples was regressed on cell type fraction estimates from InstaPrism (*θ ∈* ℝ^*N ×K*^) as follows:

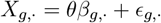

where *β*_*g,·*_ are the regression coefficients for gene *g*, and *ϵ*_*g,·*_ are the residuals.

The fraction-regressed bulk expression *X*^regress^ is then calculated by removing the predicted (fraction-attributable) component from the observed bulk and adding back the mean expression of each gene:

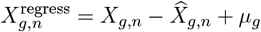

where 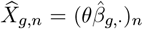 is the predicted bulk expression for sample *n*, and 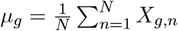is the mean expression of gene *g* across all samples.

### Top specifically expressed genes

We define *specifically expressed genes* as those differentially expressed in one cell type compared to all others. To identify these, we performed differential expression analysis at the pseudobulk level on reference scRNA-seq data: gene expression was CPM-normalized and aggregated across cells for each sample and cell type, then compared using one-versus-all tests with the `limma` R package[19]. This pseudobulk approach has been shown to enhance the robustness of marker gene detection and improve computational efficiency[20]. Genes were then ranked by the estimated log fold-change coefficients from the fitted linear model (referred to as “limma statistics” in Extended Data Figure 3), and the top *n* were selected as markers for each cell type. For our main comparison, this same set of marker genes was used for gene correlation analysis across all methods, unless otherwise specified.

## Data availability

Single-cell data used in this study were obtained from publicly available repositories or accessed with approval from the data provider. A detailed description of datasets used in this study can be found in Extended Data Table 1.

Public data: The PBMC_Perez2022 dataset[21] was accessed from the GEO platform under accession code GSE174188. The PBMC_AIDA2024 data[22] was downloaded from the the CELLx-GENE database[23] at https://cellxgene.cziscience.com/collections/ced320a1-29f3-47c1-a735-513c7084d508. The ROSMAP_AD92_Xiong2023 dataset[24] was accessed from the UCSC Cell Browser at https://cells.ucsc.edu/?ds=ad-aging-brain+ad-atac. The ROSMAP_AD430_Mathys2023 dataset[25] was downloaded from the UCSC Cell Browser at https://cells.ucsc.edu/?ds=ad-aging-brain.

The BRCA_Bassez2021 dataset[26] was downloaded from http://biokey.lambrechtslab.org. The BRCA_Wu2021 dataset[27] was downloaded through the Broad Institute Single Cell portal at https://singlecell.broadinstitute.org/single_cell/study/SCP1039. The CRC_Pelka2021 dataset[28] was obtained as a processed version from the 3CA database[29] at https://www.weizmann.ac.il/sites/3CA/colorectal under the Title identifier “Pelka et al. 2021”. The CRC_Lee2020 dataset[30] was obtained as a processed version from the 3CA database[29] at https://www.weizmann.ac.il/sites/3CA/colorectal under the Title identifier “Lee et al. 2020”. The LUAD_Kim2020 dataset[31] was downloaded from the CELLxGENE database[23, 32] at https://cellxgene.cziscience.com/collections/edb893ee-4066-4128-9aec-5eb2b03f8287 with dataset identifier “Kim_Lee_2020”. The LUAD_ Laughney2020 dataset[33] was obtained as a processed version from the 3CA database[29]at https://www.weizmann.ac.il/sites/3CA/lung under the Title identifier “Laughney et al. 2020”.

Controlled-access data: The PBMC_1k1k dataset[34] was obtained from the authors of the OneK1K Project. The ROSMAP_Mathys2024 dataset[35] was accessed from the Synapse platform under accession number Synapse ID: 52383412.

## Code availability

DALE-Eval is an open-source benchmarking framework available on GitHub at https://github.com/humengying0907/DALE-Eval. All source code, benchmarking data and detailed benchmarking results are available on Zenodo at https://doi.org/10.5281/zenodo.16649010. Both are released under the GNU General Public License v3.0.

## Acknowledgements

We gratefully acknowledge the following resources for providing data and tools used in this study: the investigators and contributors to the OneK1K Project (https://onek1k.org/); the Curated Cancer Cell Atlas (3CA) at the Weizmann Institute of Science (https://www.weizmann.ac.il/sites/3CA/); the Chan Zuckerberg Initiative Single-Cell Biology team for developing and maintaining the CellxGene platform (https://cellxgene.cziscience.com/); the Lambrechts Lab for facilitating data access (https://lambrechtslab.sites.vib.be/); the participants of the Religious Orders Study and Memory and Aging Project (ROSMAP); and the UCSC Cell Browser team for enabling interactive single-cell exploration (https://cells.ucsc.edu/).

## Author contributions

All authors contributed equally to this work.

## Competing interests

The authors declare no competing interests.

## Extended Data

**Extended Data Table 1:**
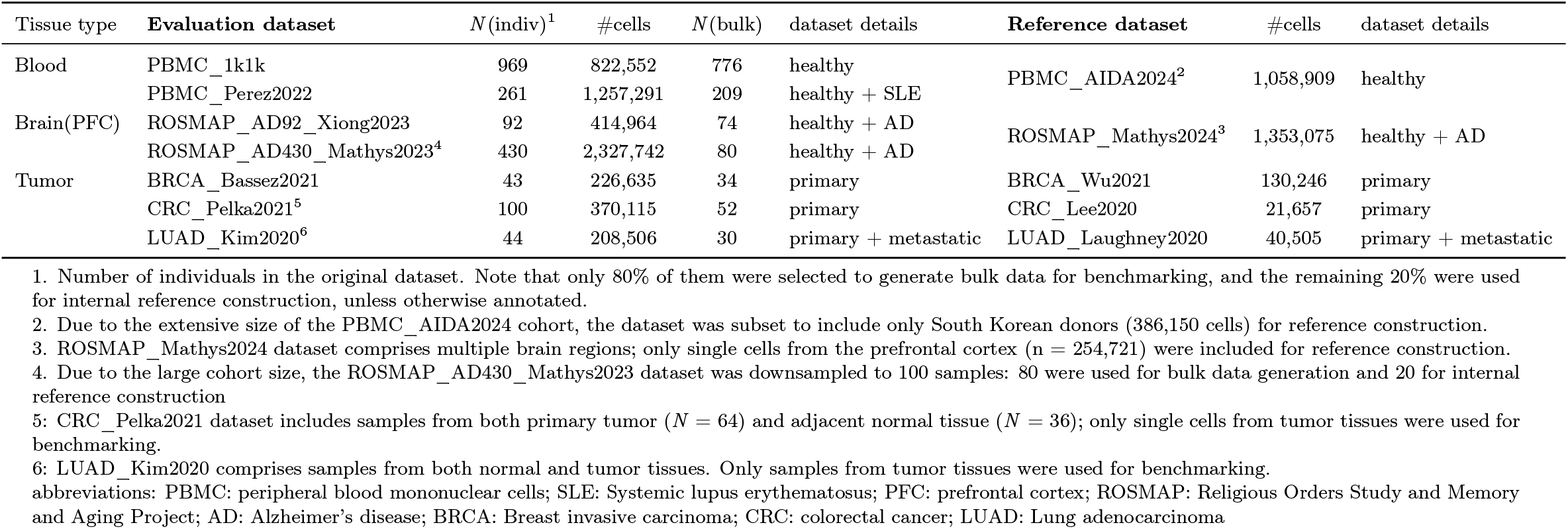
Single-cell RNA-seq dataset used for benchmarking

**Extended Data Figure 1:**
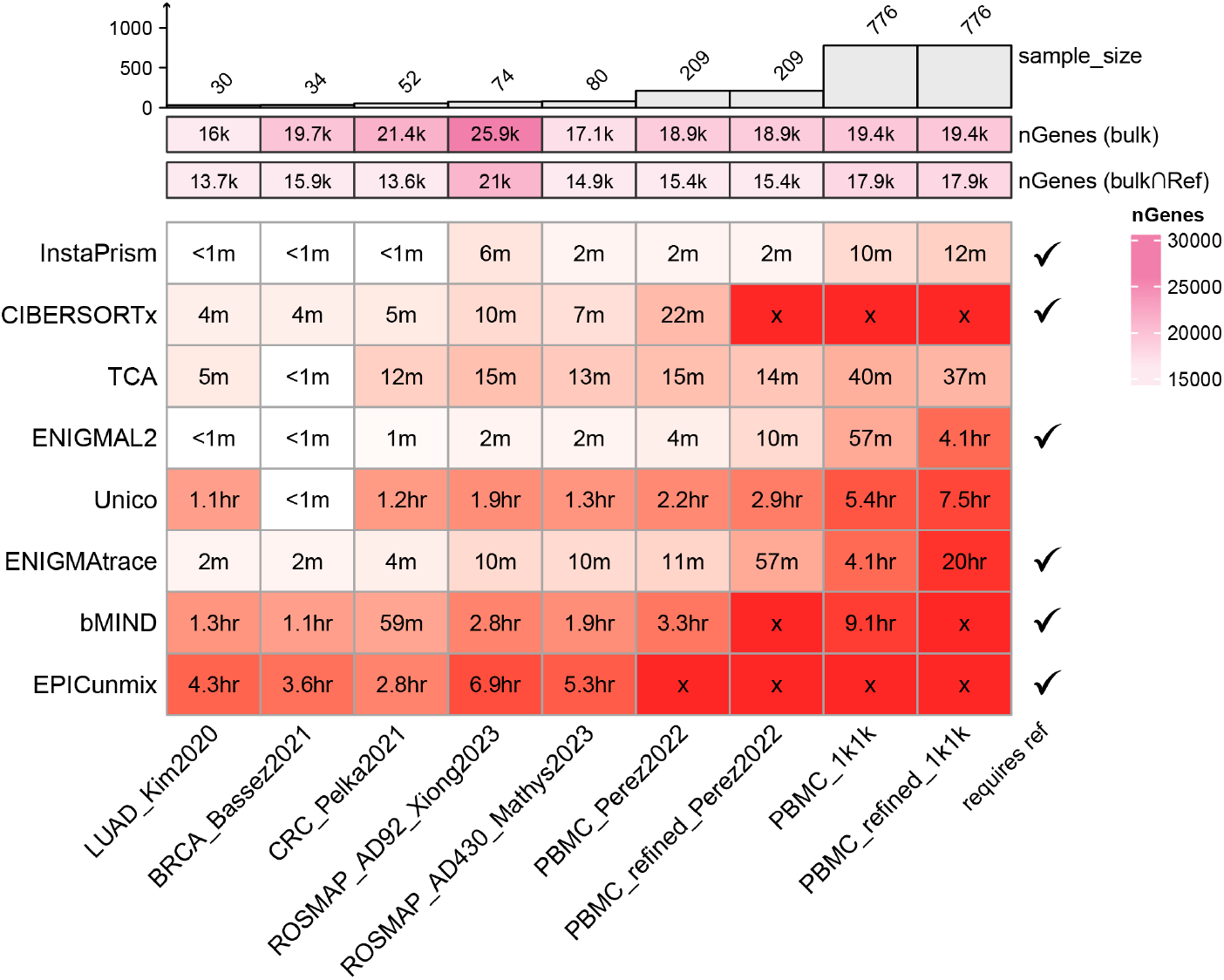
Runtime and scalability of CTSE deconvolution methods. Heatmap showing wall-clock runtime of each deconvolution method (rows) on evaluation datasets (columns); cells marked with *×* denote runs that failed to complete.

**Extended Data Figure 2:**
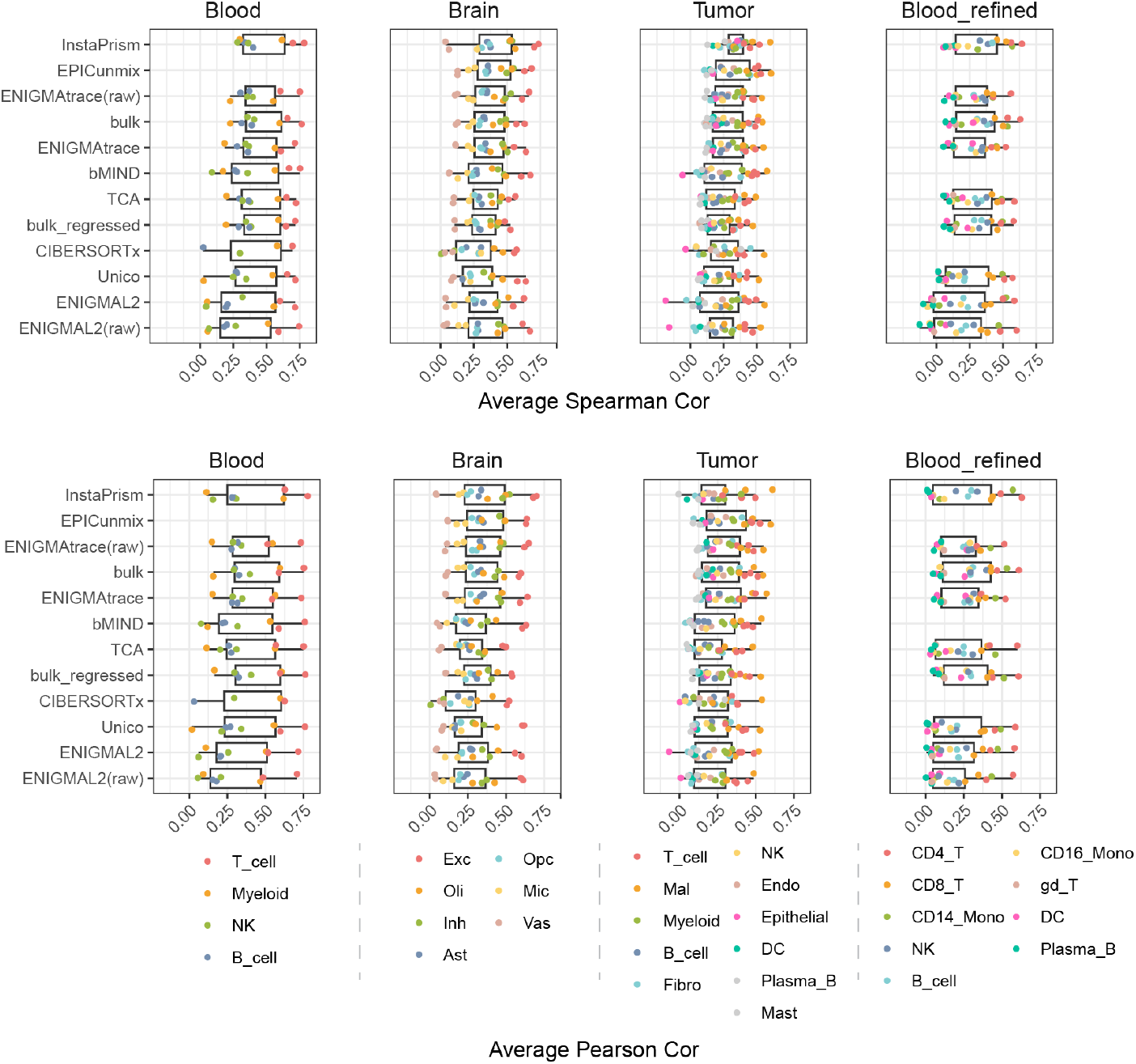
When considering all genes methods are largely indistinguishable from each other and also from bulk baselines. Boxplot showing each method’s overall deconvolution performance across tissue types: each point represents a cell type and reflects the average Spearman (top) and Pearson (bottom) correlations between ground-truth and inferred gene expression, averaged across all genes and across datasets of the same cell type within each tissue.

**Extended Data Figure 3:**
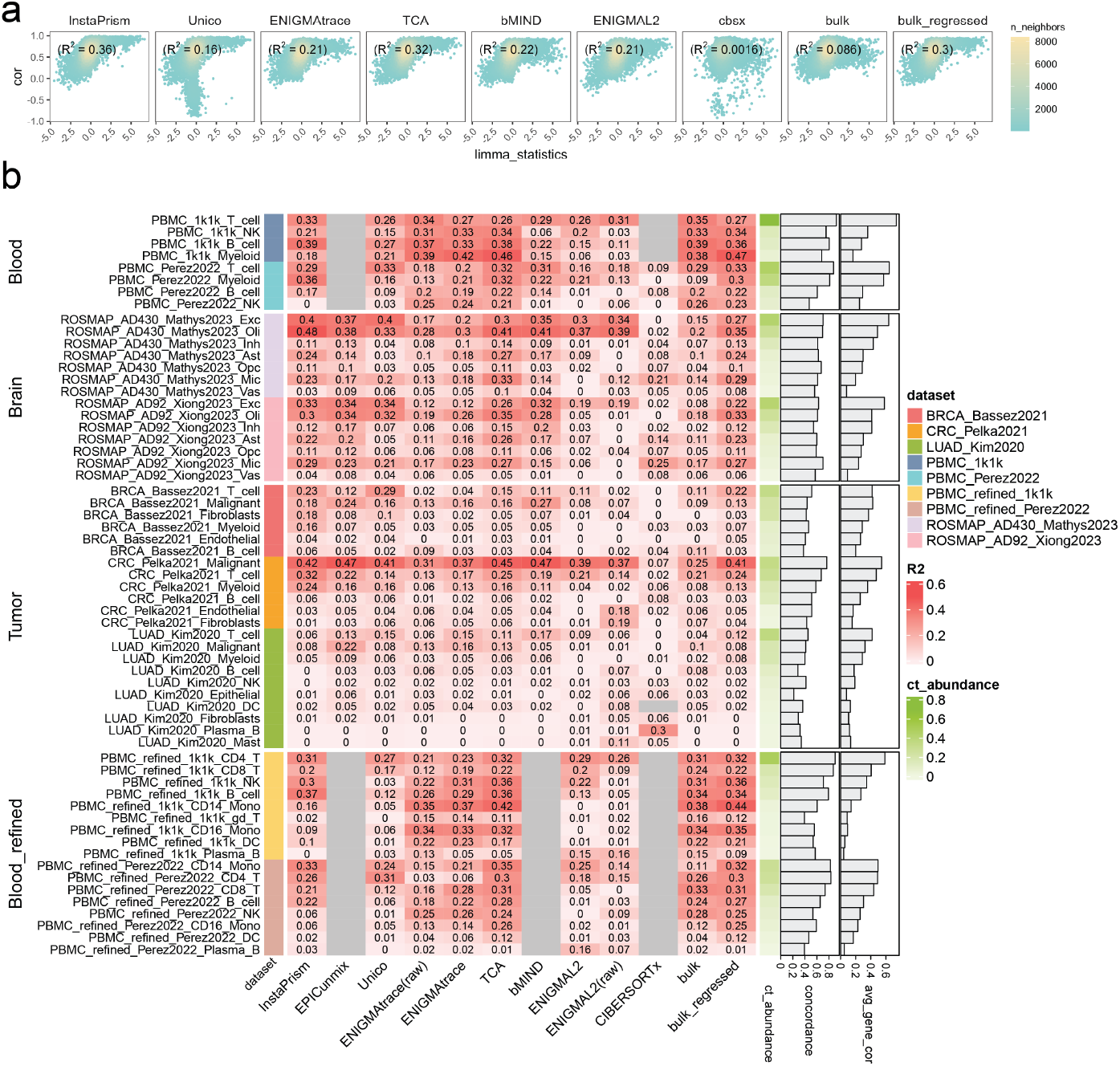
Impact of gene specificity on deconvolution performance and method concordance. **a)** Scatter plot showing how gene expression prediction accuracy (evaluated by Spearman correlation) varies with gene specificity (limma statistic). Variance explained (*R*^2^) values are annotated for each method to indicate how well specificity accounts for performance variation. Example shown for Myeloid cell from PBMC_Perez_2022 dataset. **b)**. Heatmap illustrating, for each method (rows) and cell type (columns), how gene specificity (limma statistic) explains variation in gene expression prediction accuracy (*R*^2^ values). Each cell type is further annotated by 1) its average abundance in the evaluation dataset; 2) the concordance of performance across methods, calculated as the average Pearson correlation between each method’s performance and that of the fraction-regressed bulk baseline; and 3) its average gene expression prediction accuracy for all genes, averaged across all deconvolution methods.

**Extended Data Figure 4:**
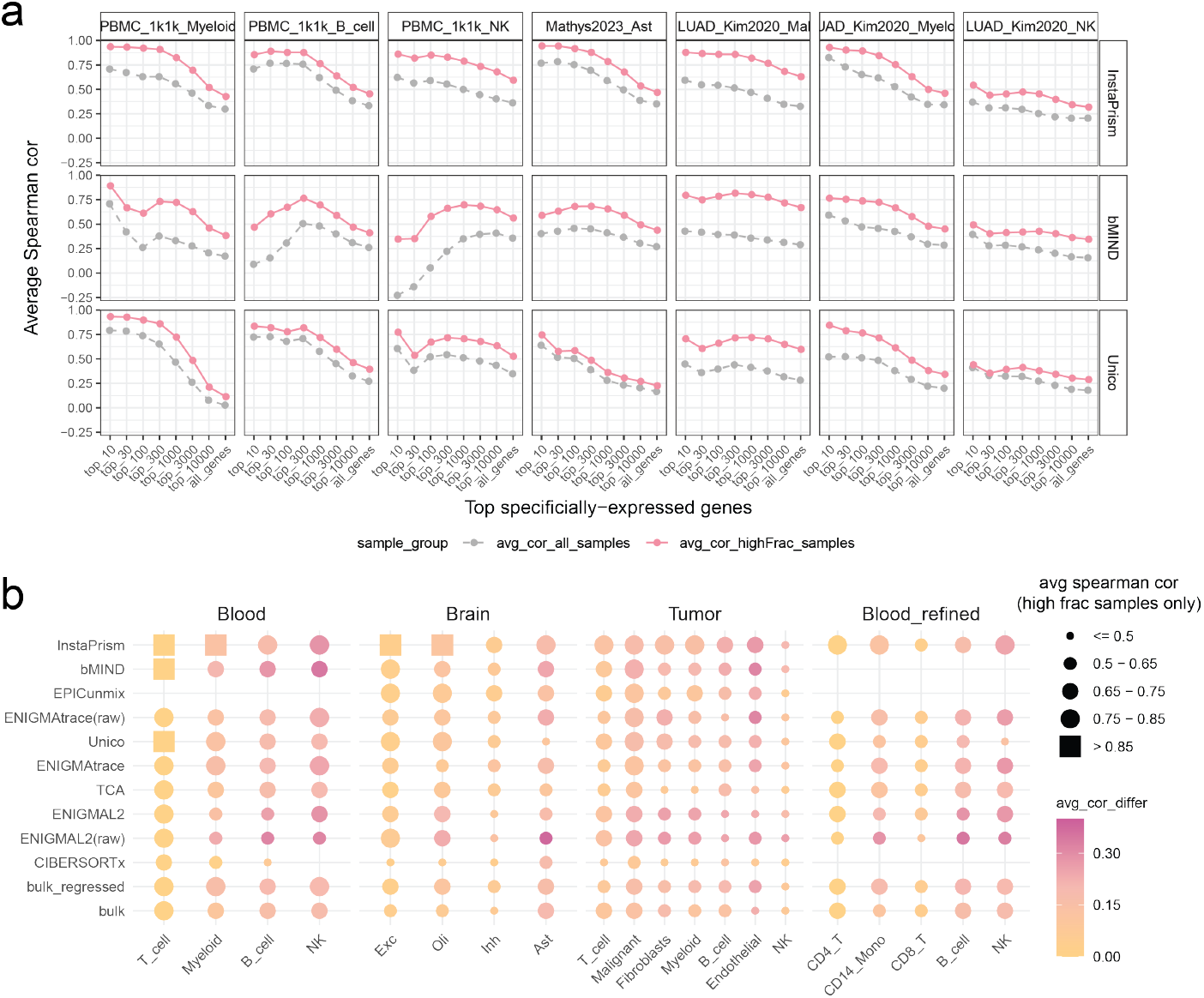
Gene correlation improvement with sample filtering. **a)** Line plot comparing gene expression prediction accuracy with and without sample filtering. Accuracy is measured as the average Spearman correlation of the top *n* specifically expressed genes. Results are shown for selected methods and datasets. **b)** Dot plot summarizing gene-expression prediction accuracy for each method *×*cell-type pair after sample filtering. Dot size corresponds to the average Spearman correlation for the top 1,000 specifically expressed genes, and color encodes the improvement achieved by sample filtering compared to using all samples. Methods are sorted by overall accuracy (baselines last), and cell types are ordered by their mean abundance after sample filtering. Methods without output are shown as empty points.

## Supplementary Information

### Supplementary Note 1: Effect of reference choices on benchmarking results

Reference profiles encode cell type–specific expression signatures for a given tissue and serve as essential inputs for CTSE deconvolution[1]. Even so-called “reference-free” deconvolution methods typically rely on known cell-type fractions, which are themselves most often estimated from a reference[2, 3]. To reflect realistic scenarios, our benchmark exclusively employed independent references, constructed from scRNA-seq datasets distinct from those used to generate the evaluation datasets. Nevertheless, we also evaluated a “self-reference” scenario, where the reference and evaluation datasets originate from the same scRNA-seq source, to benchmark the methods when the optimized reference is provided. This allowed us to benchmark method performance under optimized reference conditions and to assess if reference choice has a significant impact on benchmarking results.

Following the benchmarking framework outlined in Methods, we applied each deconvolution method to the same input data using either independent or self reference profiles. A preliminary comparison on average gene expression correlations suggested no significant differences between independent and self reference across all methods (Supplementary Figure 1a). Additionally, consistent with prior findings, gene correlation remained primarily driven by underlying cell-type abundance regardless of reference choice (Supplementary Figure 1b).

However, we note that reference choice impacts more than just signature magnitudes and exerts effects beyond the negligible differences observed in overall gene-correlation metrics. When comparing deconvolution performance in terms of fraction estimation (Supplementary Figure 1c,d), we found that using a self-reference generally resulted in higher performance, with average fraction correlation improving by approximately 0.05 compared to using an independent reference. This implies that fraction-dependent deconvolution methods may receive different input values depending on the reference choices. Consequently, future development of these CTSE methods should account for reference choice as a potential factor influencing the results, and if possible, should also assess how changes in fraction input affect downstream deconvolution outcomes.

Reference selection also alters gene prioritization. When summarizing gene correlation across the top *n* most specifically expressed genes—ranked according to different references—we observed some differences in performance results (Supplementary Figure 1e). Note that these results reflected the combined effects of differences in fraction estimates, reference input, and gene prioritization. The overall trends within each reference choice are quite similar, with generally better performance observed for more specific genes. Notably, we observed a distinct difference for the bMIND method in the myeloid cell type within the PBMC_1k1k dataset, where averaging results for the top 10 most specific genes yielded substantially different outcomes. This may be due to intrinsic instability in bMIND’s gene estimation for the most cell type-specific genes, which often display very low variation across samples.

Lastly, we compared gene correlations using the same set of top marker genes—the intersection of the top 1,000 genes ranked by self and independent reference profiles—to avoid discrepancies in gene selection (Supplementary Figure 1f). Our results suggested that while some methods performed similarly regardless of reference type, others indeed exhibited inflated performance when employing a self reference. For these methods, future improvements may lie in enhancing reference quality to more accurately capture the constituent cell-type signatures present in bulk samples.

Overall, the benchmarking conclusions remained consistent regardless of reference choice, and methods that performed well with an independent reference also ranked highly under the self-reference scenario. Our choices of independent references was able to capture trends observed with optimized references and mimics real-world deconvolution scenarios where only external references can be used[1, 4]. Future research can build upon our work by more systematically exploring how different reference selection strategies impact deconvolution performance.

**Supplementary Figure 1:**
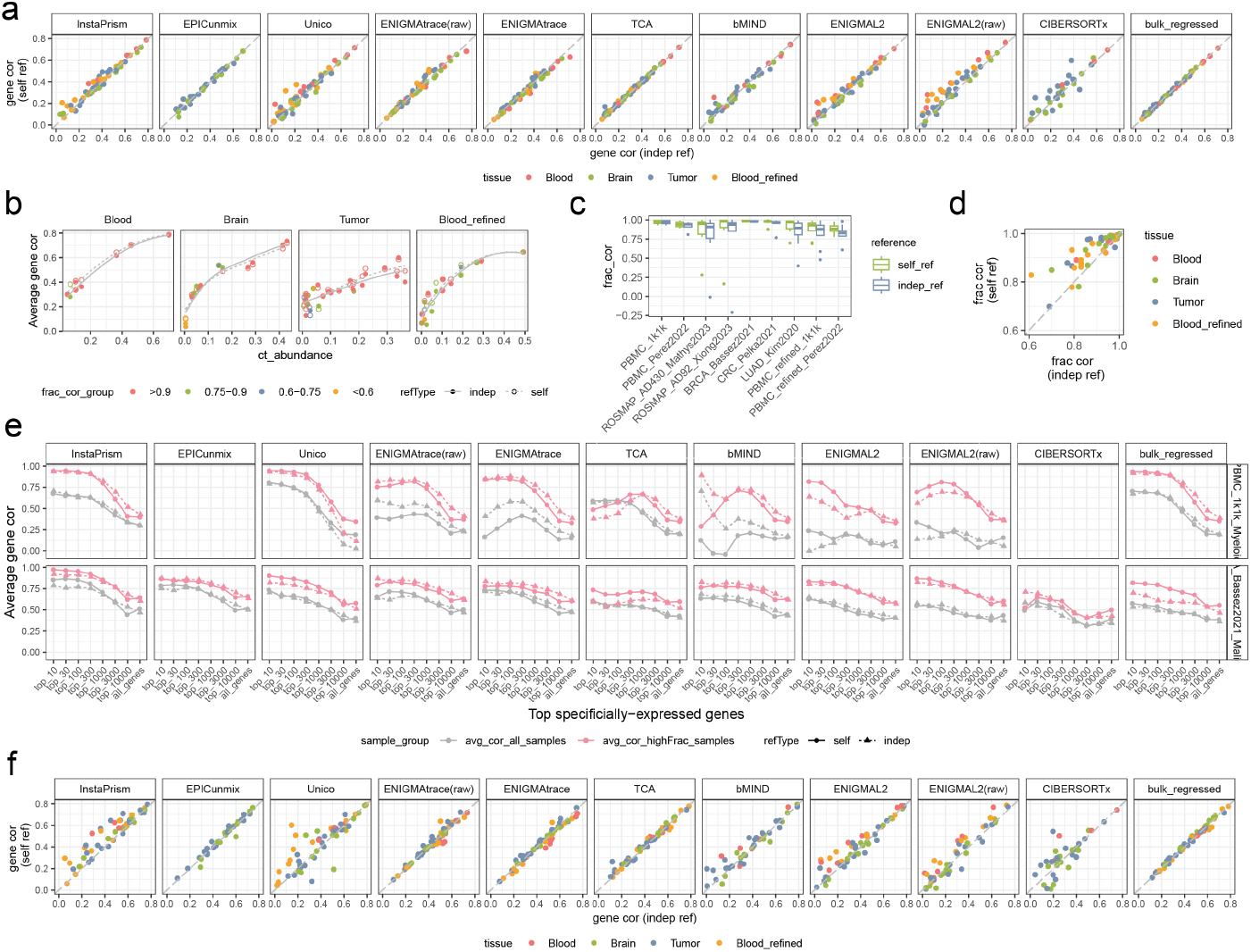
Effect of reference choices. **a)** Scatter plot comparing average deconvolution performance (mean gene expression correlation across all genes) between independent versus self references for each cell type (dot). **b)** Scatter plot showing, for both reference types, the relationship between average gene correlation and average cell type abundance. Each cell type (dot) is colored according to its fraction deconvolution performance. Results shown for InstaPrism method. **c)** Boxplot showing the distribution of fraction deconvolution accuracy for both reference choices. **d)** Scatter plot comparing fraction prediction performance between independent and self reference; cell types with fraction correlation < 0.6 are omitted. **e)** Line plot comparing gene expression correlation using self versus independent references, among the top *n* specifically expressed genes. Results are shown for selected cell types. **f)** Scatter plot comparing average deconvolution performance for the same sets of marker genes (intersection of the top 1,000 specifically expressed genes prioritized by each reference) between independent and self references.

### Supplementary Note 2. Intra vs inter sample gene correlation

This benchmark emphasizes inter-sample gene correlation as the primary evaluation metric for deconvolution performance. However, deconvolution is also frequently assessed using intra-sample gene correlation[5–7], which evaluates how well the relative expression of genes within each sample matches the ground truth. In the following section, we present results using this metric and explain why it is not considered a major basis for comparison in our study.

For this evaluation, we focus on gene correlation within hallmark genes from the MSigDB database[8], rather than using all genes. This approach highlights gene sets with well-characterized biological functions and helps avoid the inflation and bias introduced by lowly expressed genes. The resulting metric is an *N×K* matrix of correlation values, where *N* is the number of samples and *K* is the number of cell types.

Summarizing intra-sample gene correlations across all samples, most deconvolution methods achieved high accuracy, with averaging correlations > 0.85 for most cell types and clearly outperformed bulk baselines (Supplementary Figure 2a). These results are consistent with previous findings that deconvolution enables the capture of within-sample variation[5–7].

Additionally, sample-wise correlations also exhibited a clear dependence on cell-type abundance: samples with higher inferred fractions consistently demonstrated higher intra-sample correlation (Supplementary Figure 2b), suggesting that abundance estimates can help identify samples with more reliable expression profiles.

Given these seemingly promising results, we next explored whether the high intra-sample correlation observed could translate into additional biological insights, specifically through meaningful pathway scores[9–11]. For this task, we first obtained cell type-specific Meta-Program (MP) gene sets from Gavish et al.’s study[12], which previously identified expression programs defined by distinct co-variation patterns and biological functions across tumor samples. Given that these MP programs were established specifically in the context of tumor, this evaluation was conducted exclusively on tumor datasets. For each cell type-specific expression profile, we calculated pathway activity scores using Gene Set Variation Analysis[9] (GSVA, implemented via the `GSVA` R package), using gene sets from the corresponding MP gene lists. We then calculated pathway-level correlations by computing the Pearson correlation between inferred and ground-truth pathway scores.

Our results indicated that most methods failed to produce meaningful pathway variations, or did so for only a limited subset of pathways (Supplementary Figure 2c). Moreover, for most cases, pathway scores calculated from the deconvolved profiles did not even outperform bulk-derived scores.

We explored factors that might contribute to pathway-level correlations. By calculating the average inter-sample gene correlation for genes within each pathway, we observed that, in some cases—such as InstaPrism method—high pathway correlation can be explained by strong inter-sample gene correlations within that pathway. However, this relationship does not hold universally: there are cases in which gene correlation is low yet pathway scores remain high, and vice versa. Further complicating the interpretation, pathway correlations can sometimes substantially exceed the average gene correlation within the pathway. It is likely that the relative influence of gene correlation, cell-type abundance, and gene-specific weighting all contribute to observed pathway correlations. Future work should disentangle these factors to improve the accuracy and interpretability of inter-sample pathway inference.

Overall, while intra-sample gene correlation provides an additional perspective on CTSE deconvolution performance, high intra-sample correlation does not necessarily translate into biologically meaningful insights about cross-sample variation (e.g., pathway changes). Therefore, we do not consider intra-sample correlation a primary evaluation metric in our benchmark.

**Supplementary Figure 2:**
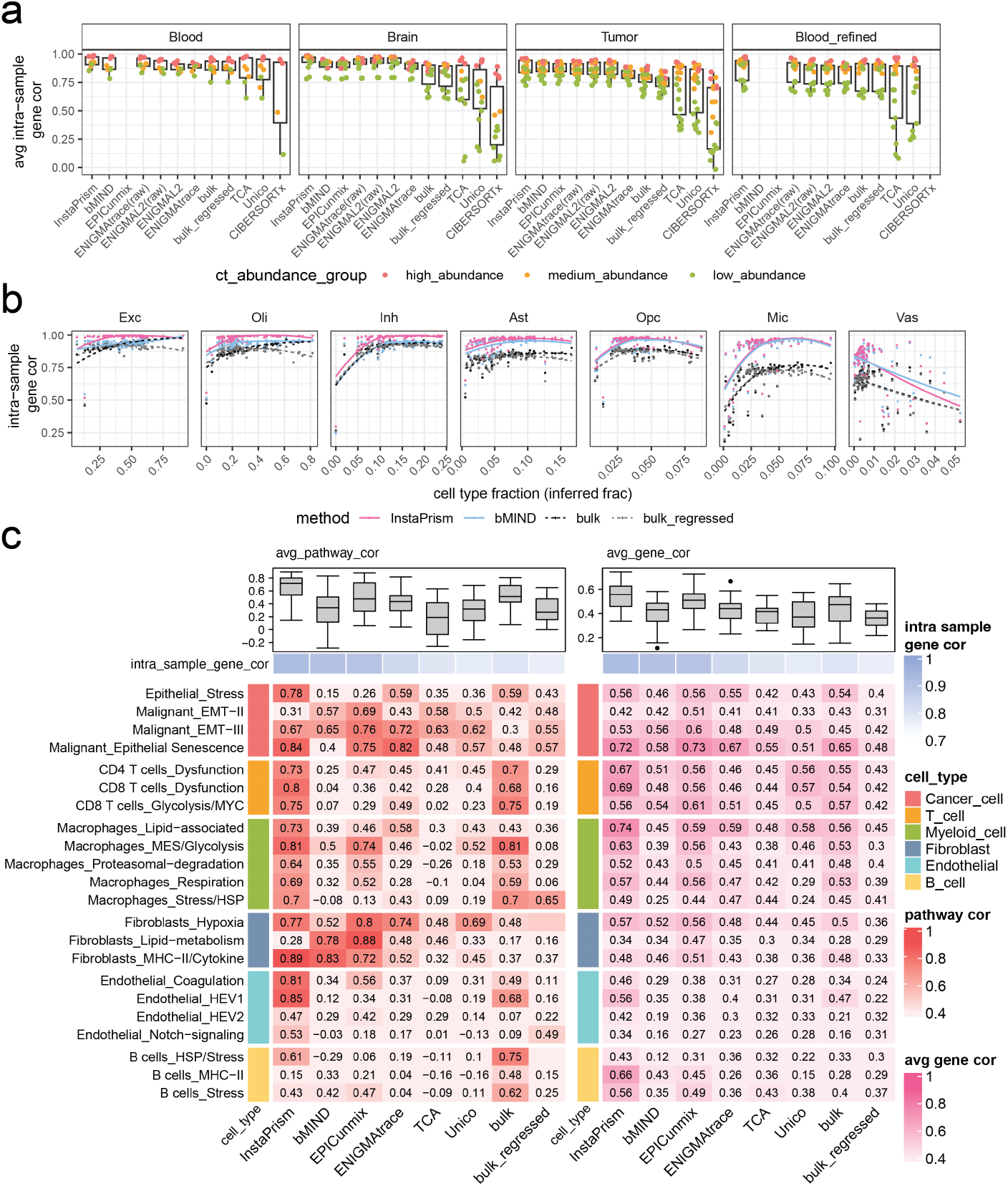
Intra-sample gene correlation and its limitation. **a)** Boxplot comparing average intra-sample gene correlation per cell type across deconvolution methods. Each cell type (point) is colored by its average abundance: high (≥ 0.25), medium (0.1–0.25), or low (< 0.1). **b** Scatter plot showing intra-sample gene correlation as a function of inferred cell-type fraction, with results shown for two top-performing methods from (a) and two bulk baselines. Example shown for dataset ROSMAP_AD430_Mathys2023. **c)** Heatmap showing (left) inter-sample pathway correlations between inferred and true pathway scores (GSVA scores) for selected cell type-specific pathways; (right) average inter-sample gene correlation across genes in each pathway. The boxplots above summarize the correlation values for each method in each evaluation setting. Example shown for BRCA_Bassez2021 dataset.

### Supplementary Note 3. Examination of cell type specificity

Cell type specificity is another key aspect of our evaluation, extending performance assessment beyond gene-level correlation alone. To illustrate this concept, we considered the expression of *CD79A*, a well-known B cell marker[13]. Examining its covariance structure in the ground-truth data revealed that variation in B cells does not correspond to variation in other cell types. However, when assessing the covariance structure in deconvolved profiles, it became immediately apparent that such variation is often propagated to other cell types as well (Supplementary Figure 3a). This phenomenon has been largely overlooked in previous studies and motivates our evaluation of covariance specificity (Methods).

On the other hand, expression magnitudes of *CD79A* varies across cell types from the CTSE profiles (Supplementary Figure 3b). Some methods, such as InstaPrism and Unico, accurately reproduced the expected pattern as seen in the ground truth for this gene, showing high expression in B cells and low expression in all others. In contrast, some methods failed to capture this specificity. For example, EPICunmix and bMIND exhibited elevated expression in non-B cell types, and ENIGMAtrace produced a B cell–specific expression distribution centered around zero. Such expression specificity is quantified using a simple metric that compares each cell type’s mean expression to the ground truth (Methods), providing a straightforward evaluation statistic for each gene (CCC value annotated in Supplementary Figure 3b).

Extending the analysis of covariance and expression specificity to other B cell markers provides a more comprehensive view of B cell specificity (Supplementary Figure 3c). For example, InstaPrism consistently exhibited lowest pairwise correlation in CTSE profiles, while methods like EPICunmix and Unico showed the most pronounced information leakage. There are also other interesting patterns: Unico sometimes displayed strong negative (inversely high) correlations; and bMIND occasionally output the same value for all B cells of a given marker, resulting in undefined and missing covariance values. Since EPICunmix builds upon bMIND results, missing covariance information in bMIND propagated to EPICunmix, leading to missing values for these genes as well. Notably, this phenomenon is not random but is most pronounced among the top marker genes for bMIND and EPICunmix. In Figure 1d, the “% of uninterpretable genes” column shows how often each method reports constant expression values across samples.

So far, our examples have focused on a few B cell–specific markers. To provide a more comprehensive view of B cell specificity, we extended the analysis to include the top 1–1000 B cell marker genes, ranked by limma-derived differential expression analysis (Methods). By computing both covariance and expression specificity for these genes, We found that specificity patterns were largely consistent across this broader gene set within each method (Supplementary Figure 3d). For the final specificity summary in Figure 2f, we averaged each cell type’s specificity scores across its top 30 specific genes to provide a representative measure of cell type specificity.

Our analysis of cell type specificity offers insights beyond basic specificity measures. The observed high cross–cell type correlations suggest that much of the variation in the deconvolved profiles does not represent biologically meaningful differences, but instead reflects signal leakage from other cell types or incomplete deconvolution. For example, gene correlations for B-cell markers within other cell types were substantially lower (Supplementary Figure 3e).

Lastly, we assessed covariance specificity from a global perspective (rather than cell type–specific) by examining the distribution of the mean off-diagonal values from the full *K× K* gene covariance matrix for each gene. This analysis revealed that all methods exhibited elevated cross–cell type covariance compared to the ground truth (Supplementary Figure 3f). The effect was most pronounced for methods such as EPICunmix and ENIGMAtrace, while relatively moderate for methods like InstaPrism and bMIND. Notably, in the LUAD_Kim_2020 dataset, cross-cell-type similarity was substantially lower, likely because the larger number of cell types—including rare populations—dilutes the average covariance values. Overall, these findings highlight the ongoing challenge of achieving truly faithful deconvolution, and suggest that future studies should further investigate strategies to improve cell type specificity.

**Supplementary Figure 3:**
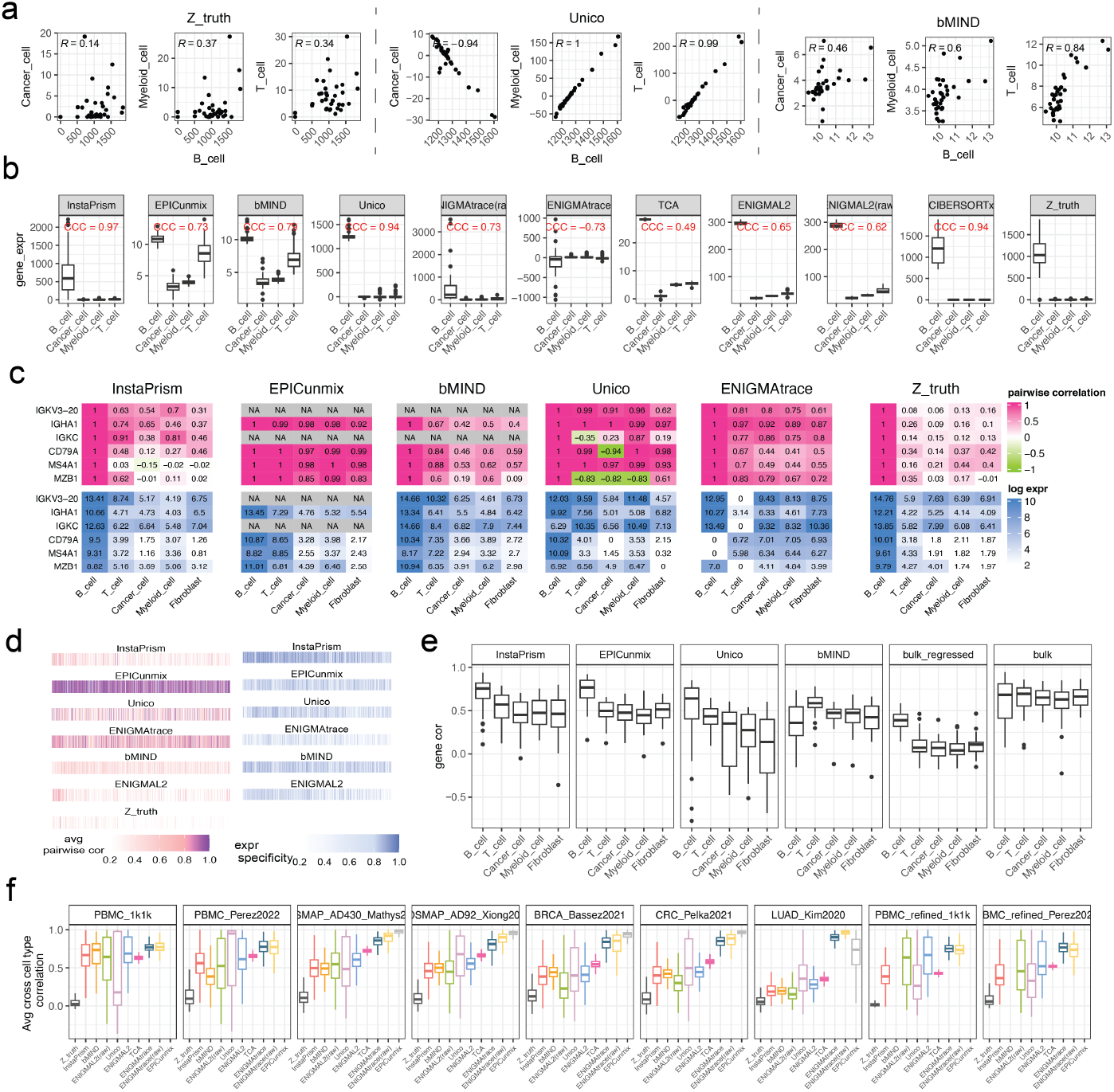
Examination of cell type specificity. **a)** Scatter plot comparing *CD79A* expression in B cell (x-axis) versus non-B cell types (y-axis) for both ground-truth and inferred CTSE; each point represents a sample. Example shown for selected methods. **b)** Boxplot showing *CD79A* expression in different CTSE profiles, labeled with expression specificity score for *CD79A*. **c)** Heatmap showing (top) pairwise correlation of B cell markers in B cells versus other cell types; (bottom) mean expression levels of these markers across all cell types (in log scale). **d** Heatmap showing pairwise correlation (pink) and expression (blue) specificity across the top 1–1,000 B cell–specific genes, ordered left to right by decreasing gene specificity. Lower pairwise correlation indicates higher covariance specificity, while higher expression scores indicate greater expression specificity. **e** Boxplot showing gene prediction accuracy for the same top 30 B cell–specific genes in B cells and in other CTSE profiles. **f)** Boxplot showing the distribution of the mean off-diagonal values from the full *K × K* gene covariance matrix, evaluated across different datasets. Panel a-e are based on results from BRCA_Bassez2021 dataset.

### Supplementary Note 4. Insights from *post hoc* analysis

We previously suggested using gene cell-type specificity as a criterion to prioritize genes likely to be accurately deconvolved (Figure 2b; Extended Data Figure 3). However, this specificity metric relies on external references, which may not fully capture marker gene specificity in the target data and could overlook certain genes. To address this limitation, we explored alternative metrics potentially predictive of gene correlation performance and thus useful for improved gene prioritization. Specifically, we considered the following metrics derived from a *post hoc* analysis of the deconvolved profiles *Z ∈* ℝ^*G×N ×K*^, each represented as a matrix of size *G × K*:

- **limma statistics:** For each cell type, we performed a one-versus-all differential expression analysis by comparing its CTSE profiles against the combined profiles of all other cell types. Specifically, CTSE profiles were first log-transformed if not already in log scale, and differential expression was assessed using the `limma` R package[14]. From the fitted linear model, we extracted the estimated log fold-change coefficients of each gene.
- **one-vs-one logFC:** As in the analysis of expression specificity, we first computed a matrix 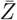 representing the mean expression of each gene in each cell type across all samples. For each cell type, we then calculated the log fold-change of the mean expression of each gene relative to the maximum mean expression observed among all other cell types.
- **partial** *R*^2^ **statistic:** For each gene, we modeled its bulk expression across samples as a multivariate linear regression on the predicted CTSE profiles of all cell types. We then extracted the partial *R*^2^ statistic for each cell type using the `rsq.partial()` function from the `rsq` R package[15], which quantifies the proportion of variance in bulk expression attributable to each cell type after adjusting for the others.

The above statistics were computed separately for each method using its deconvolution output *Z*. In addition, we also include a universal gene prioritization metric originally described in the ENIGMA study[5]:

- **GeneSigTest pvalue:** For each gene, we quantify the extent to which its variation in bulk expression is explained by the inferred cell type fractions using a linear model. We then applied Benjamini–Hochberg (BH) correction to the resulting p-values, producing an adjusted p-value matrix of dimensions *G × K*.

Applying gene prioritization using these metrics and summarizing gene correlations for genes ranked by these statistics, we observed that *post hoc* fold-change–based ranking approach offers the greatest potential for guiding gene prioritization, especially for methods with high expression specificity (Supplementary Figure 4a).

Next, we summarize deconvolution accuracy using this *post hoc* fold-change–based gene prioritization approach (Supplementary Figure 4b). For this analysis, the number of prioritized genes *n* was set proportionally to each cell type’s inferred abundance, allowing more genes to be included for cell types with higher inferred fractions. Performance was assessed only for samples with sufficient cell-type abundance (inferred fraction > 0.1). This setting represents the most optimized scenario achievable with current CTSE methods. For the top-performing method InstaPrism, we observed an average gene correlation of approximately 0.8 across all tested cell types, reflecting around a 0.1 correlation improvement over external reference-based gene ranking. However, this improvement is achieved by restricting the evaluation to highly prioritized genes and samples with adequate cell-type fractions.

Overall, this preliminary evaluation highlights the value of using a *post hoc* prioritization approach. Our *post hoc* fold-change–based ranking statistic can serve as a baseline for developing more rigorous frameworks to systematically assess confidence in gene estimates for CTSE deconvolution.

**Supplementary Figure 4:**
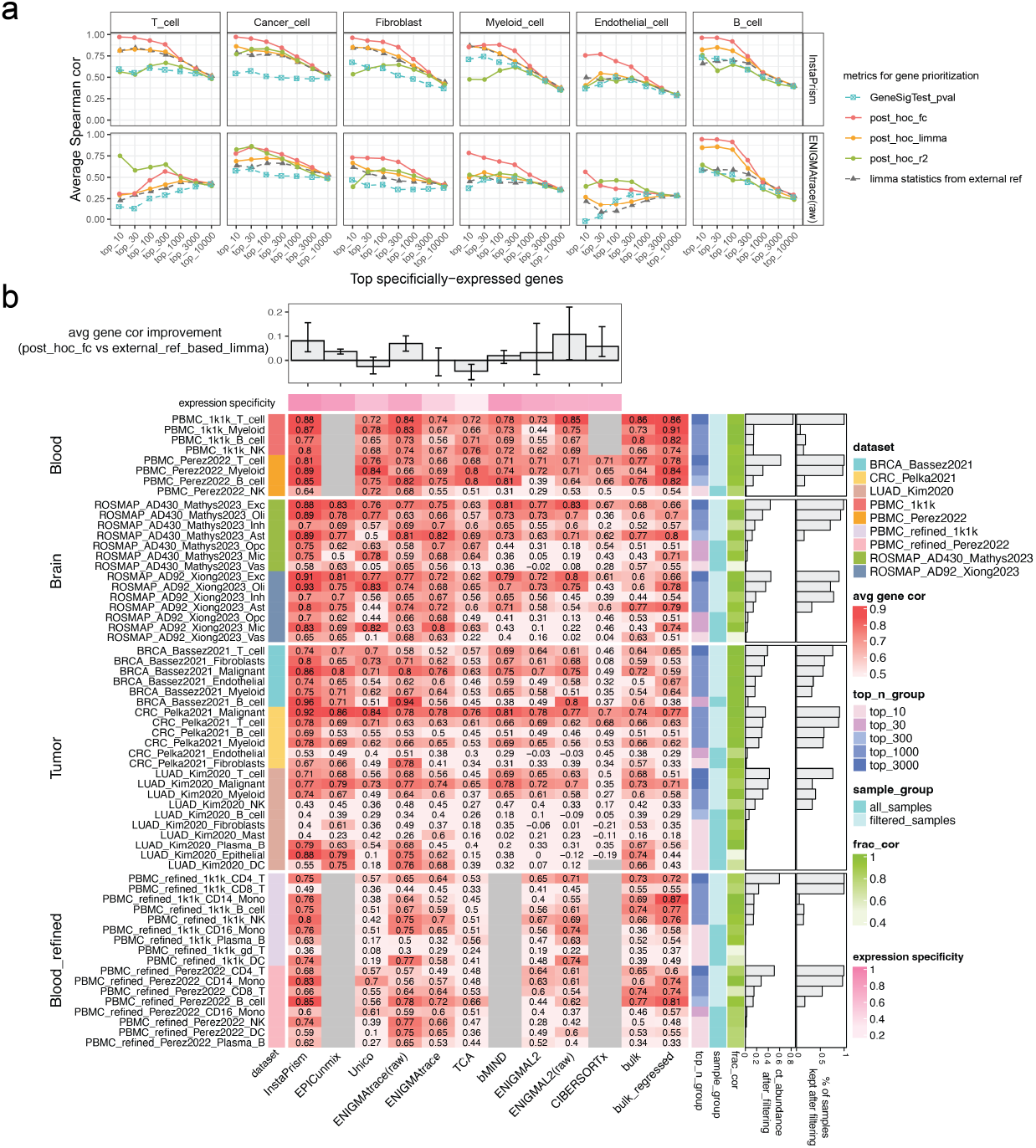
Gene correlation improvement with *post hoc* derived markers. **a** Line plot showing prediction accuracy for the top *n* specifically expressed genes, ranked by different gene prioritization methods. **b** Heatmap showing the detailed deconvolution performance of each method for the top *n* specifically expressed genes, prioritized by *post hoc* log fold change–based statistics, where *n* is scaled according to each cell type’s inferred abundance. Baseline methods use genes prioritized by reference. Spearman correlations are calculated among samples with abundance > 0.1; if fewer than 10 such samples are available, all samples are used. Each method is further annotated with its average expression specificity and the mean improvement achieved by *post hoc* log fold change-based gene ranking relative to reference-based ranking, with error bars indicating tissue-level minimum and maximum values. Each cell type is also annotated with the percentage of samples retained after filtering (abundance > 0.1) and the correlation between predicted and true fractions (frac_cor).

### Supplementary Note 5. CTSE deconvolution at higher granularity

In theory, CTSE deconvolution can resolve finer cell subtypes (e.g., CD4 and CD8 T cells) by supplying subset-level references and/or subset fractions as input. To test the feasibility of CTSE deconvolution at higher granularity, we evaluated gene correlation performance within cell subtypes (for PBMC data only). Here, ground truth was defined at greater granularity by aggregating single cells within each major cell subtype (e.g., CD4 T cells, CD8 T cells, CD14 Monocytes, and CD16 Monocytes), while the bulk input remained the same. Accordingly, both the reference profiles and the cell type fractions provided to each deconvolution method were set at the cell subtype level.

Our evaluation revealed that CTSE deconvolution becomes substantially more challenging as granularity increases. For example, for InstaPrism, the average gene correlation for T cells as a broad group was approximately 0.9, but this value dropped to around 0.7 for CD4 T cells for top 1,000 most specific genes (Supplementary Figure 5a). Additionally, our specificity analysis showed that although a gene may be highly expressed in multiple subtypes—for example, *TCRD* in gd-T[16], CD4 and CD8 T cells, the pairwise correlation between these cell subtypes is minimal inthe ground truth. However, deconvolved profiles typically display inflated covariance among these subtypes (Supplementary Figure 5b).

A detailed performance summary across methods can be found in Supplementary Figure 44b, which shows that even with the proposed optimized gene prioritization and sample filtering techniques, cell subtype–level deconvolution did not achieve the same level of reliability as observed for broad cell types. We reason that this is likely attributable to the substantial overlap of marker genes among closely related subtypes, making it significantly more challenging to deconvolve at higher resolution. Further development is needed to support high-granularity CTSE profiling with both high accuracy and cell subtype specificity.

**Supplementary Figure 5:**
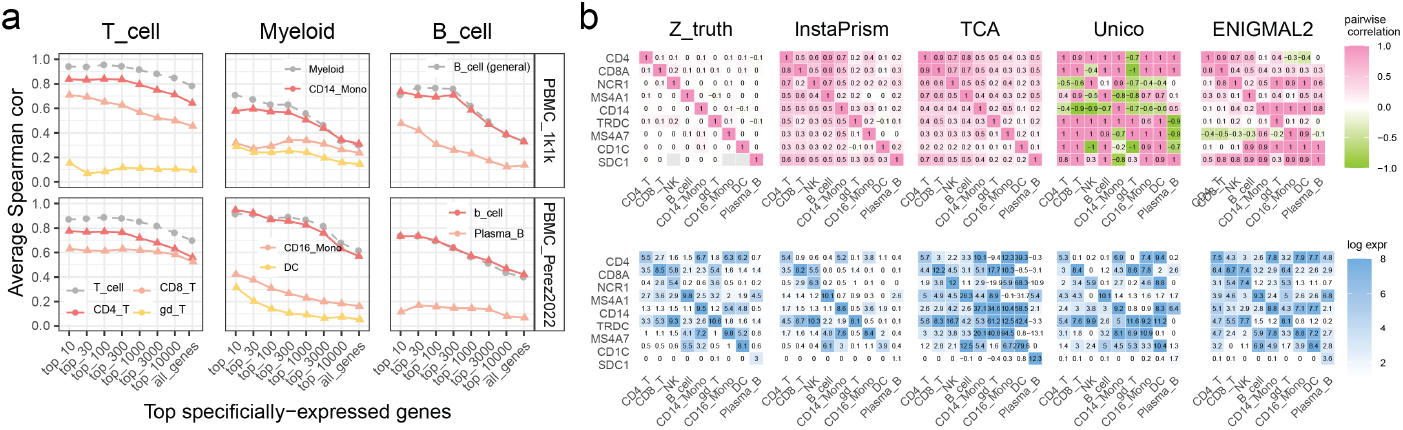
CTSE deconvolution at higher granularity. **a)** Line plot comparing gene expression prediction accuracy within CTSE profiles of different granularity, Dashed lines indicate broader cell types; solid lines represent refined cell subtypes within each broad cell type. Example shown for InstaPrism. **b)** Marker gene specificity in blood cell subtypes. Top (covariance specificity, pink): Pairwise correlation of each marker gene’s expression in its target cell type versus every other cell type. Bottom (expression specificity, blue): Mean expression of marker genes across all cell types.

## Notes

https://doi.org/10.5281/zenodo.16649010

### Competing Interest Statement

The authors have declared no competing interest.

